# Optical flow reveals motility signatures for inferring pathogenic bacterial mixture compositions via temporal convolutional networks

**DOI:** 10.64898/2026.06.29.735172

**Authors:** Yuki Fujita, Yuki Nagase, Sarthak Pathak, Alessandro Moro, Hiroaki Suzuki, Keiichiro Koiwai, Kazunori Umeda

**Author notes:** These authors jointly supervised this work (Hiroaki Suzuki, Kazunori Umeda).

## Abstract

With the rapid expansion of global food demand, aquaculture has become a critical pillar for future food security. However, aquaculture systems remain highly vulnerable to pathogenic bacteria, and rapid identification of antagonistic microbes is essential for sustainable disease control. Conventional evaluation approaches rely on fluorescence labeling or post-culture assays, limiting the ability to quantify dynamic interactions in mixed microbial populations in a real-time and label-free manner. Here, we propose a computational framework for classifying the mixing ratio of *Vibrio harveyi* and environmental bacteria using time-series motion features extracted from microscopy videos. We defined 24 interpretable motility descriptors and employed a Temporal Convolutional Network (TCN) to learn their temporal structure. The proposed method achieved a classification accuracy of 93.3%, outperforming conventional static statistical approaches and alternative machine learning models. These findings indicate that mixture discrimination in microbial communities is governed not by absolute motility magnitude, but by collective alignment and its temporal stability. Our study establishes a time-resolved computational framework for quantifying dynamic collective order in mixed microbial populations and highlights its potential for label-free automated screening and robotic microbiological applications.

**Author summary:** Bacterial infections pose a major threat to aquaculture, and rapid identification of antagonistic microbes is essential for sustainable disease management. Existing screening approaches often require fluorescent labeling or post-culture analysis, making real-time evaluation of mixed bacterial populations difficult. In this study, we show that mixture ratios of *Vibrio harveyi* and environmental bacteria can be accurately classified from time-series motion features extracted from microscopy videos. By applying TCN to 24 interpretable motility descriptors, we achieved high classification accuracy without relying on fluorescent markers. Our analysis demonstrates that collective directional alignment and its temporal stability, rather than absolute swimming speed, are the key determinants of mixture discrimination. This work introduces a computational strategy for quantifying dynamic collective order in microbial communities and supports the development of label-free, automated screening platforms for microbiological applications.

## Introduction

In recent years, global food demand has expanded rapidly due to population growth and changes in dietary patterns. According to estimates by the Food and Agriculture Organization (FAO), the world population is projected to reach approximately 9.3 billion by 2050, and food production will need to increase by nearly 60% to meet this demand [1]. Under these circumstances, aquaculture has emerged as a critical pillar for ensuring future food security. However, aquaculture systems are highly vulnerable to infections caused by pathogenic bacteria, and climate change—particularly rising sea temperatures—further increases the risk of emerging pathogens. To ensure the development of sustainable aquaculture systems, rapid pathogen detection methodologies must be established. While traditional diagnostic frameworks—including microbial culture and molecular techniques such as PCR—remain the gold standard, they are frequently constrained by significant time delays, high operational costs, and a heavy reliance on specialized laboratory infrastructure [2].

Cell identification based on imaging has been widely explored because it offers a simple and accessible approach that does not require complex molecular biological procedures or specialized analytical instruments. Such image-based methods enable rapid and non-invasive analysis, making them attractive for a variety of biological and clinical applications [3]. In recent years, there has been a notable increase in studies employing machine learning techniques to improve the accuracy and automation of cell identification, leveraging advances in computational power and data-driven modeling [4]. However, compared to eukaryotic cells, bacteria are significantly smaller, posing a fundamental challenge for image-based identification. Under conventional optical microscopy, bacterial cells often lack sufficiently distinct morphological features, making reliable classification from static images difficult [5–7]. As a result, image-based approaches for bacterial identification remain limited in their performance, highlighting the need for alternative strategies or additional sources of information beyond static imaging.

*Vibrio harveyi* is a widely distributed marine pathogen characterized by high flagellar motility and collective gene regulation via quorum sensing, making its rapid identification crucial for sustainable disease management in aquaculture [8]. Therefore, we hypothesized that analyzing time-lapse microscopy videos of microbial motility—rather than static images—would enable the identification of *V. harveyi* from diverse environmental bacterial populations without the need for fluorescent labeling or culture-based cloning.

Conventional analyses of single-cell motility predominantly rely on human-defined statistical descriptors, such as mean velocity, mean squared displacement, directional distributions, and velocity histograms [9, 10]. While these hand-crafted metrics provide useful summaries of fundamental motion characteristics [11], they inherently compress temporal information by averaging or aggregating measurements over time. As a consequence, sustained changes in motion and the sequential context of trajectories are not explicitly preserved, leading to the loss of dynamic structural information [12]. In heterogeneous microbial populations, this limitation becomes particularly critical, as the temporal dependencies and continuous evolution of motion provide essential information reflecting underlying behavioral differences. Static, pre-defined descriptors are therefore insufficient to fully capture these complex dynamics. In contrast, leveraging automated motion feature extraction allows for the direct modeling of temporal sequences, enabling the spontaneous discovery of hidden, non-intuitive motility patterns that human observers might overlook, and utilizing them as robust metrics for classification.

Furthermore, deep learning methods have recently demonstrated strong performance in image and time-series analysis [13, 14]; however, their application to microbial motility analysis remains limited. [15, 16] Convolutional neural networks (CNNs) are well-suited for extracting spatial features [14] but are not inherently designed to model long-range temporal dependencies. Recurrent neural networks (RNNs), although theoretically capable of handling sequential data, are known to suffer from gradient instability and difficulty in learning long-term dependencies [17]. Furthermore, computational frameworks that explicitly represent motility descriptors as structured time-series inputs and learn their temporal dynamics in mixed microbial systems have not been sufficiently established. Therefore, there is a clear need for computational approaches that preserve and learn the temporal structure of motility features, enabling identification of dynamically evolving collective order in mixed microbial populations.

In this study, we propose a computational framework for analyzing motility dynamics in mixed microbial populations while preserving temporal structure. Specifically, we extract motion features of cell populations from time-series microscopy images using optical flow. This representation captures velocity fields and their spatiotemporal variations, including directional tendencies, spatial organization, and temporal fluctuations of collective motion. By transforming raw image sequences into a motion-based representation, our approach enables analysis within a physically interpretable feature space, facilitating the interpretation of learned representations in terms of biologically meaningful motility characteristics. We then employ a Temporal Convolutional Network (TCN) [18] to learn temporal patterns in the extracted motion features. TCN [18] leverages dilated causal convolutions, enabling efficient modeling of long-range temporal dependencies while maintaining stable training dynamics. By operating on motion representations derived from optical flow, the model captures subtle temporal signatures of bacterial motility that are not readily discernible from individual frames or by human observation. As a result, our approach enables the estimation of the relative abundance of *V. harveyi* within heterogeneous bacterial populations from microscopy image sequences. Although the present study focuses on estimating the abundance of *V. harveyi* in mixed populations, the proposed framework is generalizable and has the potential to be extended to the detection and characterization of other bacterial species.

## Methods

### Dataset

In this study, we analyzed images of bacterial samples consisting of the marine pathogen *V. harveyi* and environmental bacteria. *V. harveyi* was provided by Prof. K. Koiwai [19]. Environmental bacteria were collected from soil sampled from a flower bed at the Korakuen Campus of Chuo University (Bunkyo, Tokyo, Japan). The bacteria were isolated through dilution, centrifugation, and filtration. Both bacterial strains were stored as glycerol stocks in a freezer. For experimental use, a small aliquot was inoculated into a culture medium (2.5% Brain Heart Infusion medium; Becton & Dickinson, Cat#211065, supplemented with 1.5% NaCl) and incubated overnight with shaking at 25?C. The cell concentrations of both bacterial populations were then estimated from microscopy images and diluted to the desired concentration (5 × 10^8^ cells/mL). The mixing ratio between *V. harveyi* and environmental bacteria was systematically varied from 0:10 to 10:0 in increments of 0.5, resulting in 21 distinct mixture conditions. For each mixture ratio, independent mixing and imaging experiments were conducted.

Microscopic observations were performed using an inverted microscope (Nikon Ti [20]). Time-lapse images were recorded with a CCD camera under the bright-field conditions (Fig 1). Each video was captured at a spatial resolution of 1920 × 1200 pixels. For each sample, 27 consecutive frames were acquired at a frame rate of 9 fps. For each mixture condition, imaging was repeated 10 times while varying the field of view in order to account for spatial heterogeneity and diverse distribution patterns of bacterial populations. To reduce spatial bias and increase the number of independent training samples, each 1920 × 1200 frame was subdivided into six non-overlapping regions of 640 × 600 pixels. This preprocessing step generated six sub-sequences from each original images. Across all mixture conditions, a total of 1,260 time-series samples were constructed (21 mixture ratios with 60 samples per condition). For deep learning analysis, the dataset was divided into training and testing subsets. For each mixture condition, 50 samples were randomly selected for training, and the remaining 10 samples were reserved for testing. This balanced sampling strategy ensured an equal number of samples across mixture ratios, preventing bias toward specific conditions and enabling fair evaluation of mixture classification performance.

**Fig 1.**
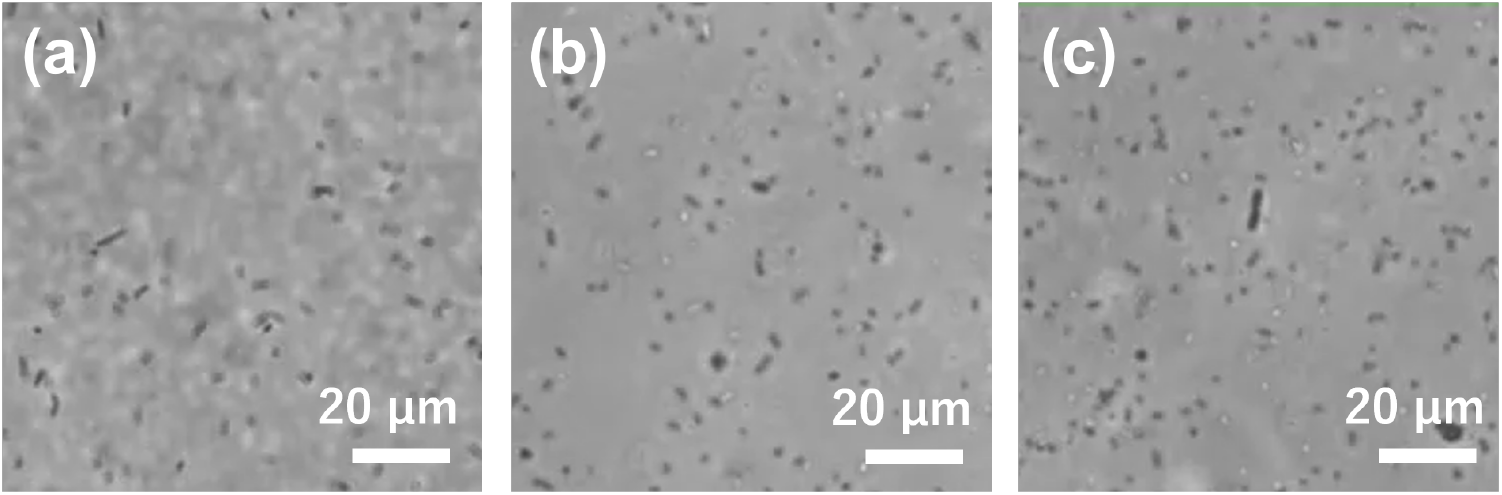
Snapshots of original images with mixing ratios of (a) 10:0, (b) 5:5, and (c) 0:10 (*V. harveyi* : environmental bacteria).

### Motion Feature Extraction

In the following, we describe the proposed motion feature extraction framework, illustrating each step with concrete examples from the experimental data. Pixel-wise motion between consecutive frames was estimated using the dense optical flow method proposed by Farnebäck [21]. This approach approximates local image neighborhoods with quadratic polynomials and estimates displacement by analyzing changes in polynomial coefficients across frames, enabling dense estimation of subtle and continuous motion. In microscopy videos, both individual-level micro-movements and population-level collective flows can coexist; thus, obtaining dense motion information at the pixel level is essential. Fig 1 shows snapshots from the original consecutive images at three different mixing ratios of *V. harveyi* and environmental bacteria. The bacteria appear as black dots or rods, making it virtually impossible to identify their species from still images alone. From the estimated flow field (*u, v*), we extracted motility descriptors to represent collective dynamics in a low-dimensional yet interpretable manner. In this study, the features were grouped into the following six categories: (i) velocity statistics, (ii) directional statistics, (iii) motion diversity indices, (iv) frequency-domain features, (v) spatial gradient and autocorrelation indices, and (vi) local motion pattern indices. These categories were designed to quantify collective motion from complementary perspectives, including motion intensity, directional order, distributional complexity, temporal fluctuation structure, spatial continuity, and local deformation patterns. The 24 motility features used in this study were designed to systematically and comprehensively characterize the motion state of bacterial populations from a physical perspective. The above six categories—covering velocity, direction, temporal variation, and spatial order—were organized by referring to physically meaningful quantities commonly used in active matter research and collective motion analysis (e.g., order parameters and alignment statistics) [22–24], as well as standard frequency-domain and statistical measures widely used in signal processing [25]. Therefore, the proposed feature space is not an empirical collection of arbitrary statistics but a structured representation grounded in theoretical frameworks of collective motion.

#### Velocity statistics

Velocity statistics were computed from the magnitude of the optical flow to quantify the overall motility and velocity distribution of bacterial motion. Let *u* and *v* denote the horizontal and vertical components of the optical flow, respectively. The flow magnitude at each pixel is defined as

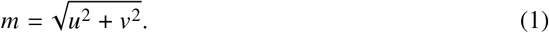

Using the magnitude values of all pixels in a frame, several statistical measures were computed to characterize the distribution of motion intensity. First, the mean flow magnitude,

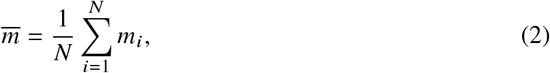

represents the average activity level of bacterial motion across the entire image. To capture the heterogeneity of motion intensity, the standard deviation of the flow magnitude was calculated as

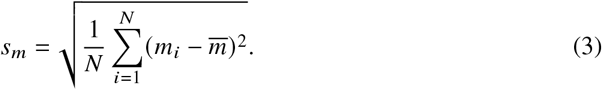

This quantity reflects the degree to which fast and slow movements coexist within the observed population. To obtain a robust representation of the distribution, the median value of the flow magnitude,

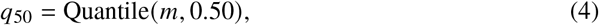

was used as a representative statistic that is less sensitive to extreme values.

In addition, higher-order quantiles were computed *q*_75_, *q*_90_, *q*_95_, *q*_99_, which capture the tail structure of the velocity distribution. These statistics are particularly useful for detecting localized regions exhibiting strong motion. To further analyze the characteristics of high-velocity regions, pixel sets corresponding to the top 25%, 10%, 5%, and 1% of flow magnitudes were defined as *S*_25_, *S*_10_, *S*_5_, and _1_, respectively. The average velocity within each subset was calculated as

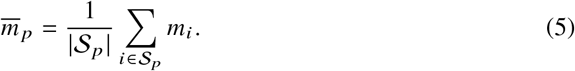

This measure enables the detection of highly motile regions even when the overall average motion level is low. Fig 2 shows an example of the spatial distribution of the flow magnitude. To evaluate the spatial extent of active motion, the proportion of moving pixels was also calculated. Let 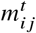 denote the flow magnitude at pixel (*i, j* ) between frame *t* and *t* + 1. A motion threshold *τ*^*t*^ was defined as

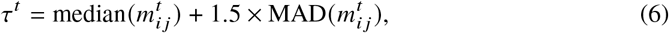

where MAD denotes the median absolute deviation, a robust statistic against outliers. Given an image of height *H* and width *W*, the proportion of pixels exceeding the threshold is computed as

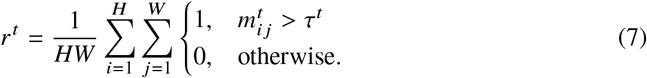

**Fig 2.**
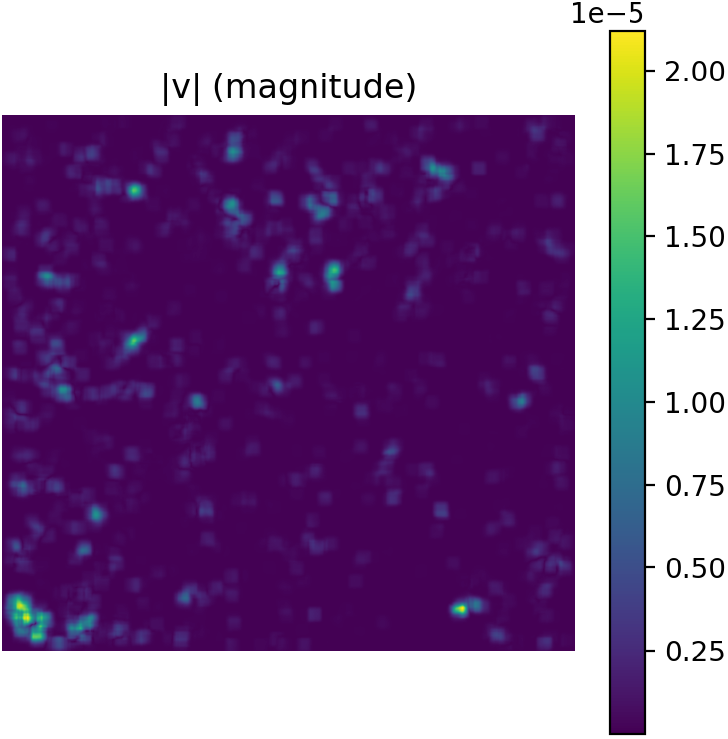
Magnitude of the velocity vector for each pixel at a mixing ratio of 5:5.

Finally, the average proportion of moving pixels over the entire video sequence was calculated as

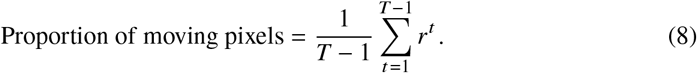

Unlike simple velocity averaging, this measure evaluates the spatial prevalence of fast-moving regions and is therefore effective for detecting localized populations of highly motile bacteria. An example of the motion mask for the mixture ratio 5:5 is shown in Fig 3.

**Fig 3.**
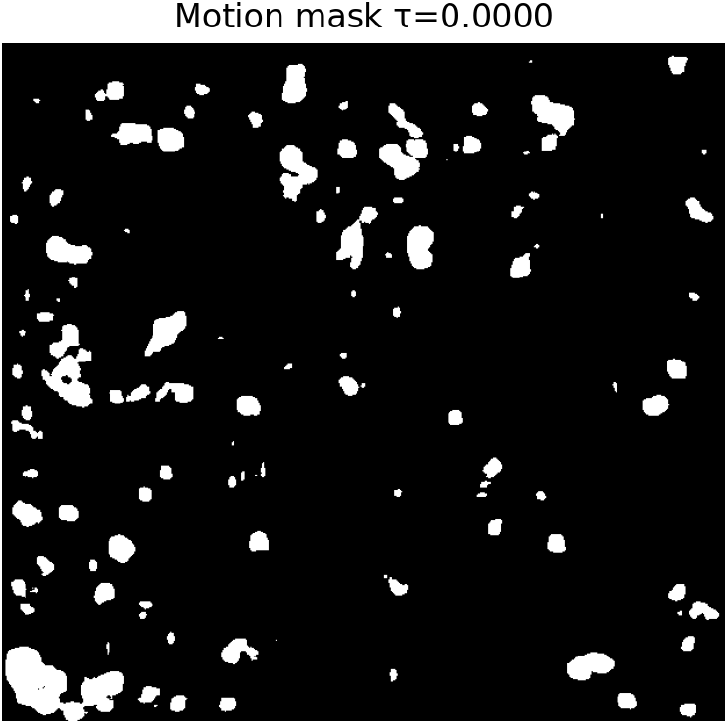
Mask image of moving pixels with a mixing ratio of 5:5.

#### Directional statistics

Directional statistics were used to quantify the degree of alignment and randomness in the motion directions of bacterial populations. The direction of motion at each pixel was computed from the optical flow components as

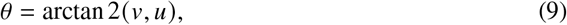

where *u* and *v* denote the horizontal and vertical components of the flow vector, respectively. To characterize the dispersion of motion directions, the entropy of the directional histogram was calculated as

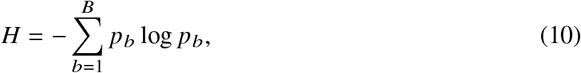

where *p*_*b*_ represents the probability of directions falling into the *b*-th bin of the histogram with *B* bins. To capture temporal characteristics of directional variability, the mean entropy and its temporal fluctuation were computed as

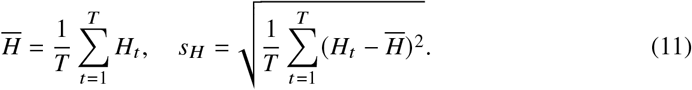

These quantities enable the distinction between motion patterns in which directions remain consistently aligned and those in which directional disorder varies over time. In addition to entropy-based measures, the concentration of motion directions was evaluated using the circular mean vector defined as

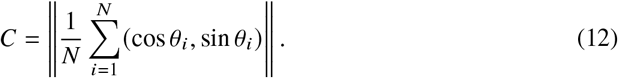

The temporal mean and temporal variation of this quantity, denoted by 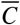 and *s*_*C*_, were also calculated. These measures quantify the degree of collective alignment in the motion field: higher values indicate coherent motion toward a common direction, whereas lower values correspond to more dispersed or disordered motion patterns.

#### Motion diversity indices

Motion diversity indices were introduced to quantify the complexity of motion intensity and directional distributions within the bacterial population. Since simple statistical measures such as the mean or variance are often insufficient to capture differences in distribution shapes, entropy-based measures were employed to evaluate the diversity of motion. Specifically, the entropy of the motion magnitude distribution was calculated as

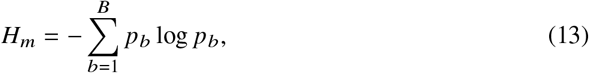

where *p*_*b*_ represents the probability of motion magnitudes falling into the *b*-th bin of a histogram with *B* bins. This measure allows the distinction between motion fields concentrated around a single velocity level and those composed of multiple distinct motion components. In other words, it characterizes the complexity and irregularity of the motion field. Higher entropy values indicate that diverse motion magnitudes and directions coexist within the population, whereas lower values suggest more uniform and ordered motion patterns. Thus, this index provides a quantitative criterion for distinguishing between relatively homogeneous motion patterns and more diverse collective behaviors. Fig 4 shows a color visualization of the flow field in the HSV color space, where hue represents the motion direction and brightness represents the magnitude of velocity. Fig 5 shows a polar histogram of motion directions for the mixture ratio 5:5. These visualizations provide an intuitive representation of the distribution of motion magnitude and direction in the bacterial population.

**Fig 4.**
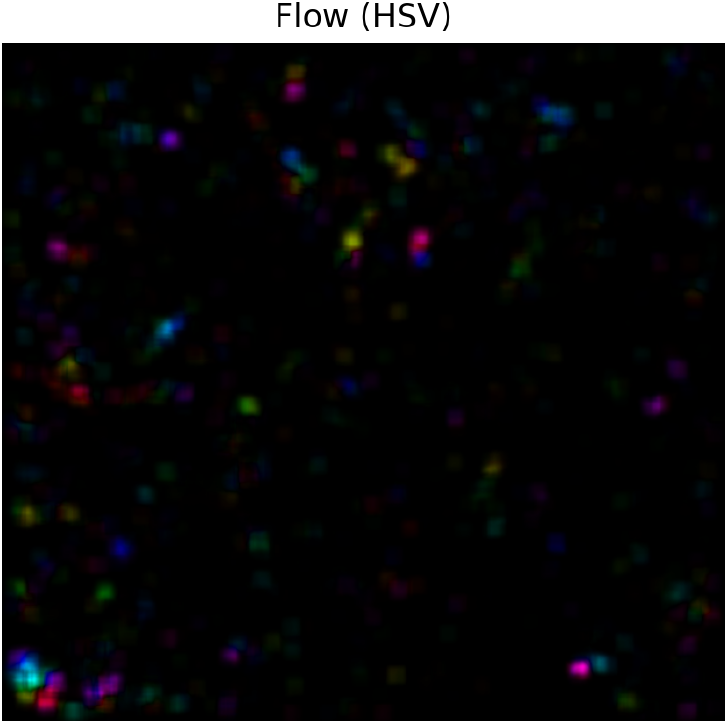
HSV color visualization chart of direction and magnitude of velocity.

**Fig 5.**
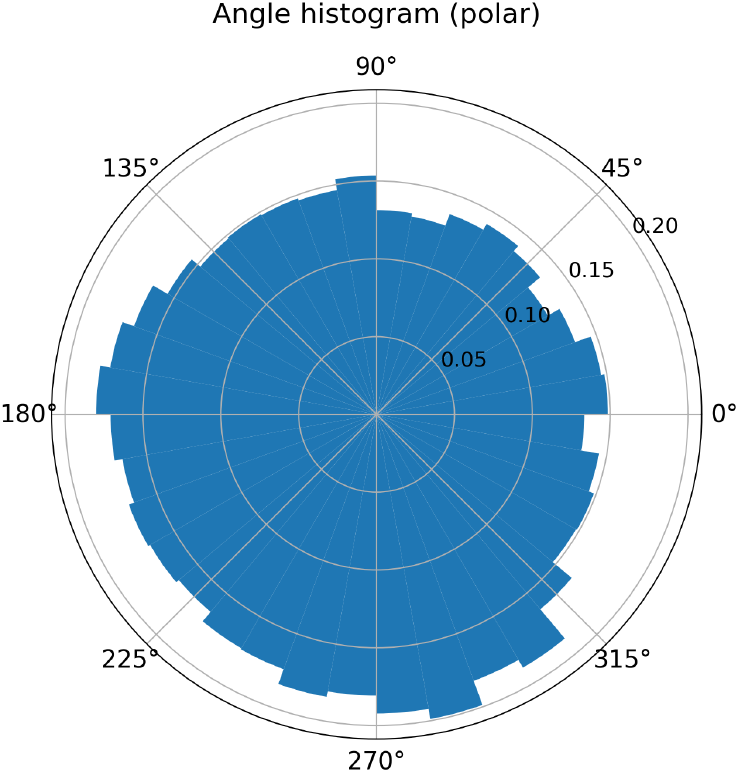
Polar coordinate histogram of angular distribution for a mixing ratio of 5:5.

#### Frequency-Domain Features

Frequency-domain features were introduced to characterize temporal variations in the motion field, including periodic fluctuations and fine-scale temporal dynamics. In this study, power spectrum analysis was performed by applying the Fast Fourier Transform (FFT) to a temporal scalar sequence derived from frame-to-frame motion information. Given a temporal sequence *x*_*t*_ (*t* = 0, 1, …, *T* − 1), the discrete Fourier transform was computed as

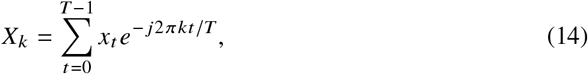

where *X*_*k*_ represents the complex Fourier coefficient for frequency index *k*. The corresponding power spectrum was then obtained as

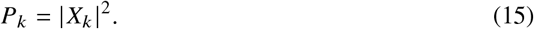

To separate large-scale smooth motion from fine and rapid fluctuations, the frequency spectrum was divided into low-frequency and high-frequency components. A cutoff radius was defined in the frequency domain to separate the low-frequency set ℒ and high-frequency set ℋ. The proportion of high-frequency components was computed as

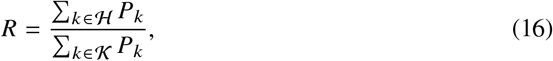

where *K* denotes the set of all frequency components. To capture temporal characteristics of the motion dynamics, the temporal mean and temporal standard deviation of this ratio were calculated as 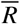, *s*_*R*_. A higher value of the high-frequency ratio indicates that rapid and fine-scale motion fluctuations dominate the motion field, whereas lower values suggest smoother and more coherent large-scale motion. This feature therefore quantifies the balance between local rapid movements and globally smooth motion patterns. Fig 6 shows the logarithmic power spectrum obtained from the FFT of the frame-difference images. The cutoff circle used to separate the low-frequency and high-frequency regions is also illustrated.

**Fig 6.**
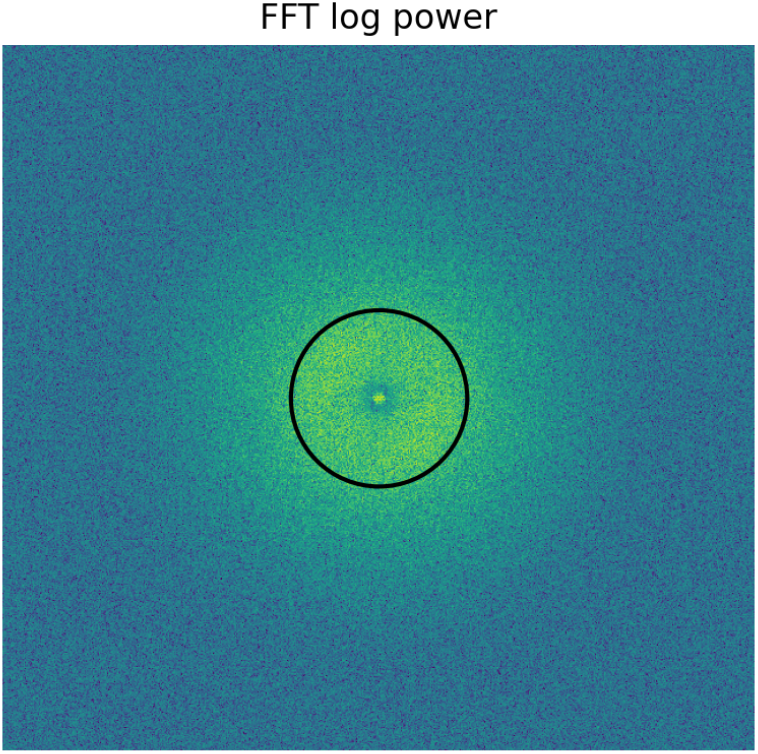
FFT log power spectrum and cutoff circle of difference images with a mixing ratio of 5:5.

#### Spatial gradient and autocorrelation indices

Spatial gradient and autocorrelation indices were introduced to quantify the spatial continuity and granularity of the motion field. These features characterize whether motion is smoothly distributed over space or composed of locally heterogeneous structures. The magnitude of the spatial gradient was computed as

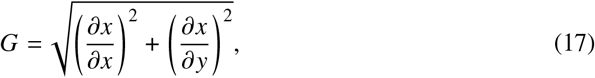

which measures the rate of spatial change in the motion field. Large gradient values indicate abrupt spatial variations, while smaller values correspond to smoother motion distributions. To capture temporal characteristics of the spatial variation, the temporal mean and temporal standard deviation of the gradient magnitude were calculated as 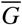, *s*_*G*_. In addition, spatial autocorrelation was computed to quantify the degree of spatial coherence in the motion field. The autocorrelation function was defined as

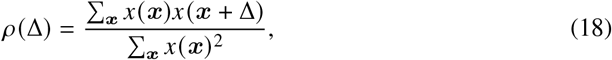

where Δ denotes the spatial displacement vector and ***x*** represents the spatial coordinates. The mean autocorrelation value 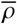 was used as a summary statistic. These measures reflect the spatial organization of the motion field. High autocorrelation values indicate spatially coherent motion that extends over larger regions, whereas low values correspond to fragmented motion patterns with strong local variations. Thus, these features enable discrimination between smooth collective motion and locally irregular motion structures.

#### Local motion pattern indices

Local motion pattern indices were introduced to characterize localized structures and deformation patterns in the motion field. These features capture spatial flow behaviors such as expansion, contraction, rotation, and the formation of coherent motion domains.

First, divergence and vorticity were computed from the spatial derivatives of the optical flow field. The divergence of the flow field was defined as

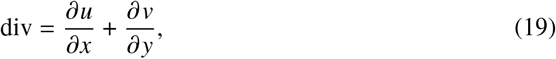

where *u* and *v* denote the horizontal and vertical components of the flow vector. The mean and standard deviation of the divergence values were calculated to quantify the degree of local expansion and contraction. Positive divergence indicates locally divergent motion, where the flow spreads outward, whereas negative divergence indicates convergent motion. Fig 7 shows an example divergence map for the mixture ratio 5:5.

**Fig 7.**
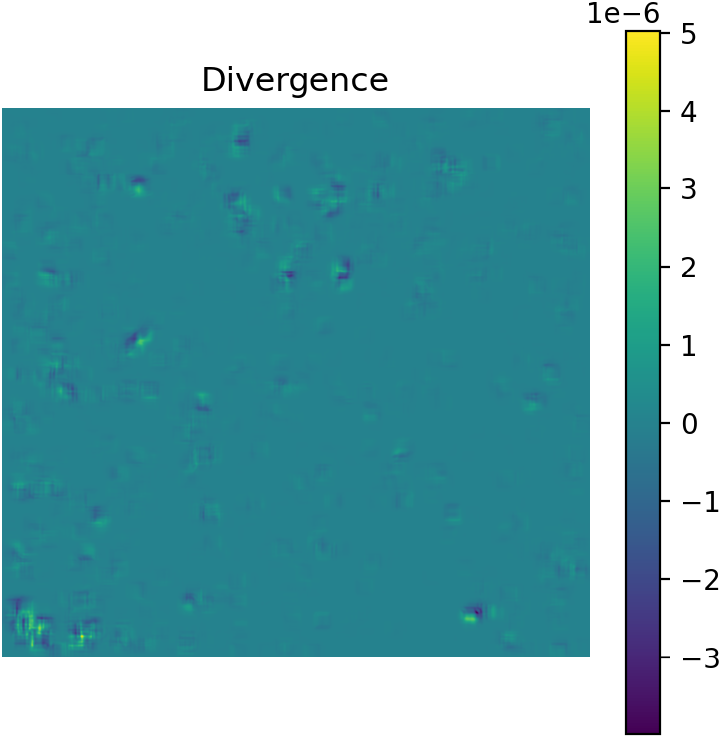
Divergence map with a mixing ratio of 5:5.

Next, local rotational motion was quantified using the vorticity defined as

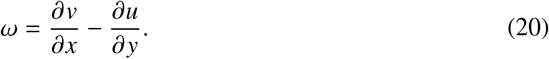

The mean and standard deviation of the vorticity were computed to evaluate the strength of rotational structures in the motion field. Larger vorticity values indicate the presence of strong rotational flow patterns, whereas smaller values correspond to primarily translational or diffusive motion. An example vorticity map for the mixture ratio 5:5 is shown in Fig 8.

**Fig 8.**
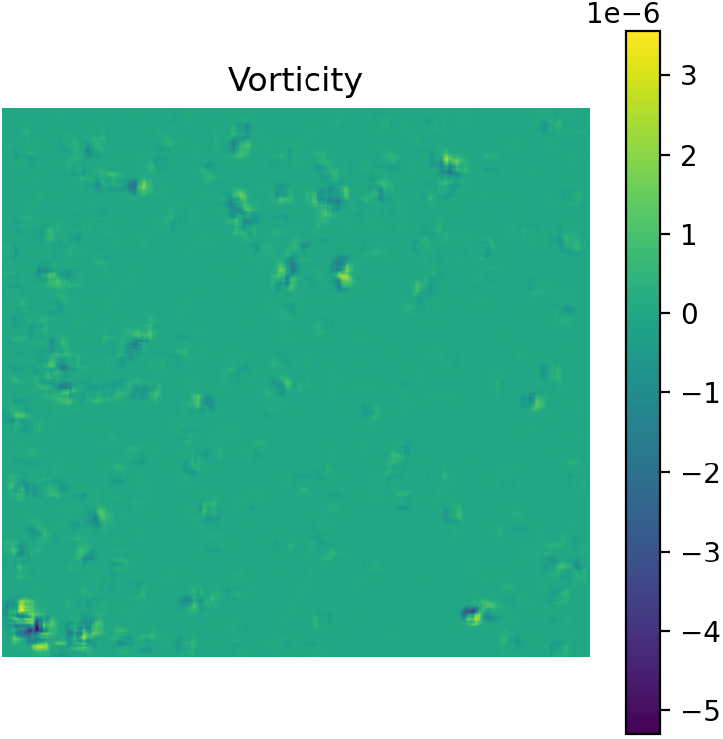
Vorticity map for a mixing ratio of 5:5.

In addition, coherent motion regions were extracted by binarizing the flow magnitude and performing connected-component analysis. Let *K*_*t*_ denote the number of connected motion components in frame *t*. The temporal mean of the component count was calculated as

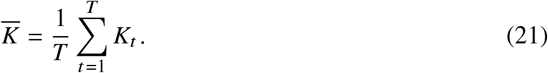

For each connected component, the area was computed, and the mean and standard deviation of the component areas were obtained as 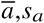,*s*_*a*_. Furthermore, the 95th percentile of the component area distribution, *q*_95_ *a*, was used to emphasize the presence of large motion domains. These indices characterize whether motion occurs as numerous small fragmented regions or as large coherent clusters. Consequently, they enable discrimination between dispersed motion patterns and coordinated collective motion structures.

### Temporal Convolutional Network

To classify bacterial mixture ratios from the extracted motion feature sequences, we employed a TCN [18]. TCN models temporal dependencies using one-dimensional convolutions and can efficiently capture long-range temporal relationships through dilated convolutions [18, 26]. Compared with RNNs, TCNs allow parallel computation and are less susceptible to gradient vanishing or exploding problems. Fig 9 illustrates the overall processing flow of the proposed TCN-based classification workflow. The input to the network is a time series of 24-dimensional motion features. Each sample consists of 27 frames, and the input sequence is represented as

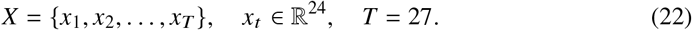

**Fig 9.**
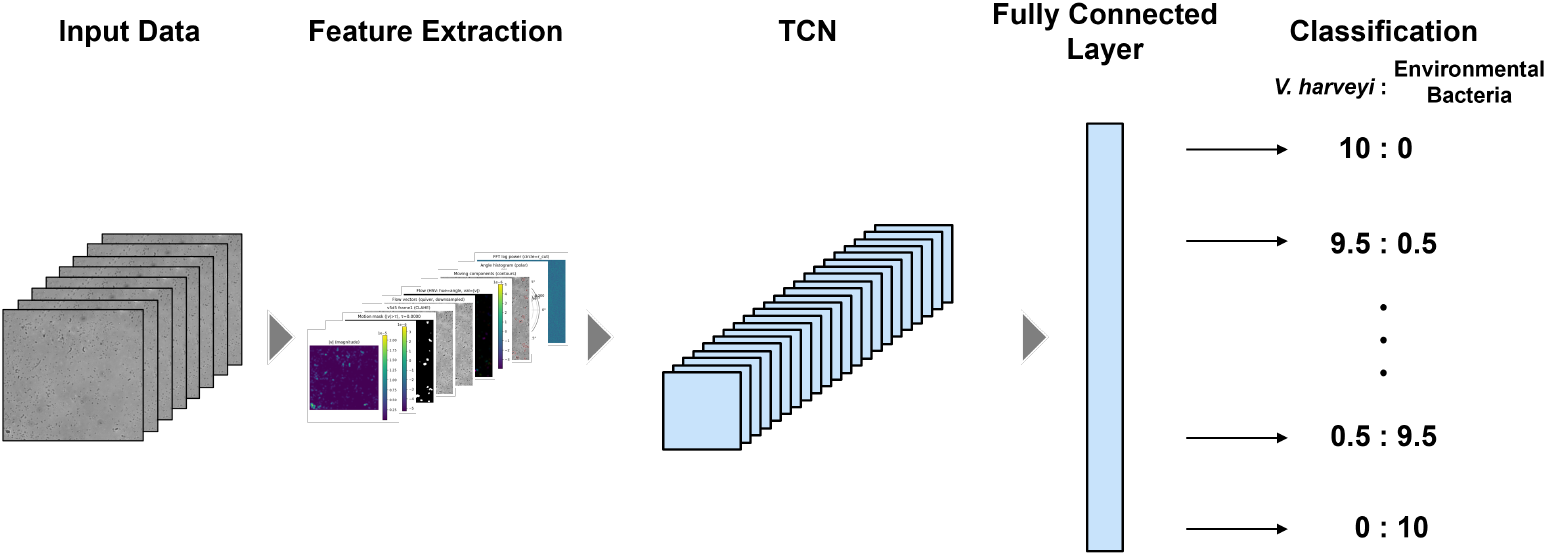
Flow diagram of the motion feature extraction and TCN-based classification workflow.

The first layer of the network is a 1 × 1 convolutional projection layer that maps the 24 input features to a 128-dimensional channel space. This layer learns linear combinations of the motion descriptors while providing a higher-dimensional representation suitable for subsequent convolutional processing. The projection layer is followed by three residual TCN blocks. Each block contains two dilated convolution layers with batch normalization, ReLU activation, and dropout, together with a residual connection. Let *h*^(*l*)^ denote the input to the *l*-th block and *F*(·) the nonlinear transformation within the block. The residual output is expressed as

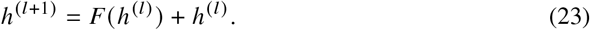

The residual connections stabilize gradient propagation and improve optimization in deeper architectures. To capture temporal dependencies at multiple scales, dilated convolutions with increasing dilation factors were employed. In this study, the dilation factors were set to *d* = 1, 2, 4. With a kernel size of *K* = 3, this configuration yields an effective receptive field of approximately 15 frames. This enables the model to capture both short-term fluctuations and mid-range temporal structures in bacterial collective motion. The temporal feature representation produced by the TCN [18] blocks was aggregated using global max pooling along the time dimension. Max pooling emphasizes the most prominent motion patterns observed within each sequence. Finally, the pooled representation was passed to a fully connected layer that outputs classification scores for the 21 mixture-ratio classes. The model was trained using the cross-entropy loss

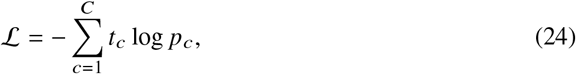

where *t*_*c*_ denotes the ground-truth label and *p*_*c*_ the predicted probability for class *c*. In this study, *C* = 21 corresponds to the number of mixture conditions. Model parameters were optimized using the AdamW optimizer. The main hyperparameters included a kernel size of 3, 128 channels, dropout rate of 0.2, learning rate of 0.002, and batch size of 8. These settings allow the model to learn both short-term fluctuations and the temporal evolution of collective motion patterns.

### Comparative Models

To evaluate the effectiveness of the proposed TCN [18], we conducted comparative experiments using several machine learning models. The baseline methods included Support Vector Machine (SVM) [27], a one-dimensional convolutional neural network (1D-CNN) [28], and XGBoost [29]. For the SVM [27] model, the time-series motion features were first converted into fixed-length feature vectors using temporal statistics. Specifically, the mean, standard deviation, quartiles (25%, 50%, and 75%), and maximum value were computed for each feature channel across time. These statistics were concatenated to form a fixed-length representation for each sample. The resulting vectors were then used as inputs to an SVM [27] classifier with a radial basis function (RBF) kernel. SVM [27] performs maximum-margin classification in a high-dimensional feature space and is widely used as a robust baseline model for nonlinear classification problems. To compare with deep learning approaches, 1D-CNN [28] was also implemented. The input consisted of the 24-dimensional motion feature sequences. First, a 1 × 1 convolution layer was used to project the feature channels into a higher-dimensional representation. Subsequently, one-dimensional convolutional layers were applied to capture local temporal patterns in the motion features. The resulting feature maps were aggregated using global max pooling along the temporal dimension and fed into a fully connected layer to perform mixture-ratio classification. 1D-CNNs are widely used for modeling sequential data and have been successfully applied to time-series analysis.

In addition, we evaluated XGBoost [29], a gradient-boosted decision tree algorithm designed for efficient learning from tabular data. Unlike TCN [18] and 1D-CNN [28], XGBoost [29] does not directly model temporal order; instead, it learns nonlinear decision boundaries from aggregated motion descriptors. In this study, the extracted 24 motion features were used as input for multi-class classification. The hyperparameters of XGBoost [29] were determined by five-fold cross-validation, and the final model was trained using the selected configuration. The optimized hyperparameters were a maximum tree depth of 5, 800 boosting trees, a learning rate of 0.05, a subsampling ratio of 0.8, a feature subsampling ratio of 0.8, an *L*_2_ regularization term *λ* = 3.0, and a minimum child weight of 1.0. These settings were adopted for the final XGBoost [29] model used in the comparative experiments.

All models were trained and evaluated using the same dataset splits and feature representations to ensure fair comparison.

### Feature Importance Analysis

To analyze which motion features contributed most to the classification of bacterial mixture ratios, we conducted a feature importance analysis using permutation importance. Permutation importance evaluates the contribution of each feature by measuring the decrease in model performance when the relationship between that feature and the target variable is disrupted. In this study, the importance of each motion feature was evaluated using the trained TCN [18]. Specifically, the motion feature sequences consisted of 24 feature channels extracted from optical flow fields. For each feature channel, the temporal order of the feature values within each sample was randomly shuffled while keeping all other features unchanged. This operation destroys the temporal structure of the selected feature while preserving its statistical distribution.

The permuted feature sequences were then input into the trained TCN [18] model, and the classification accuracy was recalculated using the test dataset. The importance score of each feature was defined as the decrease in classification accuracy caused by the permutation. To reduce the influence of randomness, the permutation procedure was repeated multiple times for each feature channel, and the mean and standard deviation of the accuracy drop were computed. In this study, five repetitions were performed for each feature. Features that produced larger decreases in classification accuracy were interpreted as having greater contributions to the classification of bacterial mixture ratios. The resulting permutation importance values provide insights into which physical aspects of bacterial collective motion are most informative for mixture-ratio estimation.

## Results

### Overall Classification Performance

To evaluate the effectiveness of the proposed approach, the extracted 24 motion features were input to TCN [18] to classify the mixture ratios of *V. harveyi* and environmental bacteria. The resulting confusion matrix is shown in Fig 10. The proposed model achieved an overall classification accuracy of 93.3%, demonstrating that the mixture ratios can be reliably identified from the motion features.

**Fig 10.**
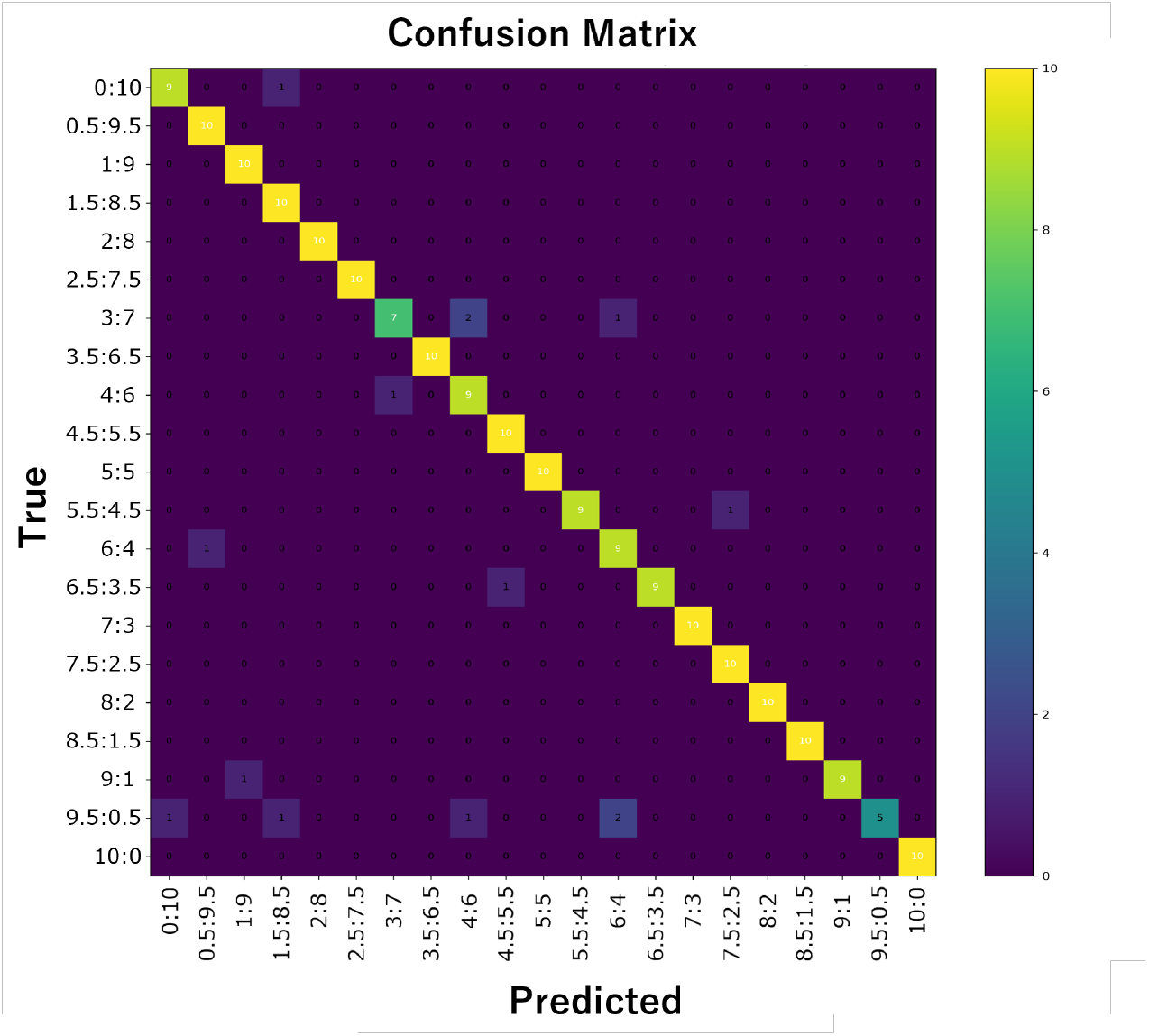
Confusion matrix of mixture-ratio classification using the 24 motion features with TCN.

To further analyze the performance across individual classes, precision, recall, and F1 score were calculated for each mixture condition. These metrics are defined as follows. Precision is defined as the ratio of true positives to all samples predicted as positive for a given class, expressed as Precision = TP/(TP + FP), where TP denotes true positives and FP denotes false positives. A high precision value indicates that when the model predicts a sample as belonging to a particular mixture-ratio class, this prediction is likely to be correct, meaning the model generates few false alarms. Recall is defined as the ratio of true positives to all samples that actually belong to a given class, expressed as Recall = TP/(TP+FN), where FN denotes false negatives. A high recall value indicates that the model successfully identifies the majority of samples belonging to a particular mixture-ratio class, meaning few true instances are missed. The F1 score is the harmonic mean of precision and recall, defined as F1 = 2 · (Precision × Recall)/ (Precision + Recall). This metric provides a balanced evaluation that accounts for both false positives and false negatives, and is particularly informative when the performance trade-off between precision and recall needs to be summarized in a single value. The results are summarized in Fig 11, Fig 12, and Fig 13.

**Fig 11.**
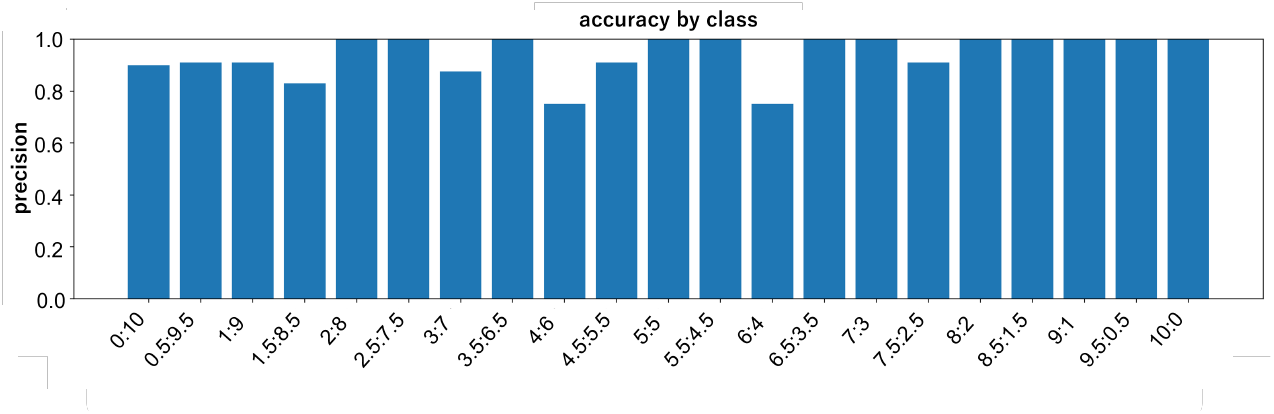
Class-wise accuracy of mixture-ratio classification using the 24 motion features with TCN.

**Fig 12.**
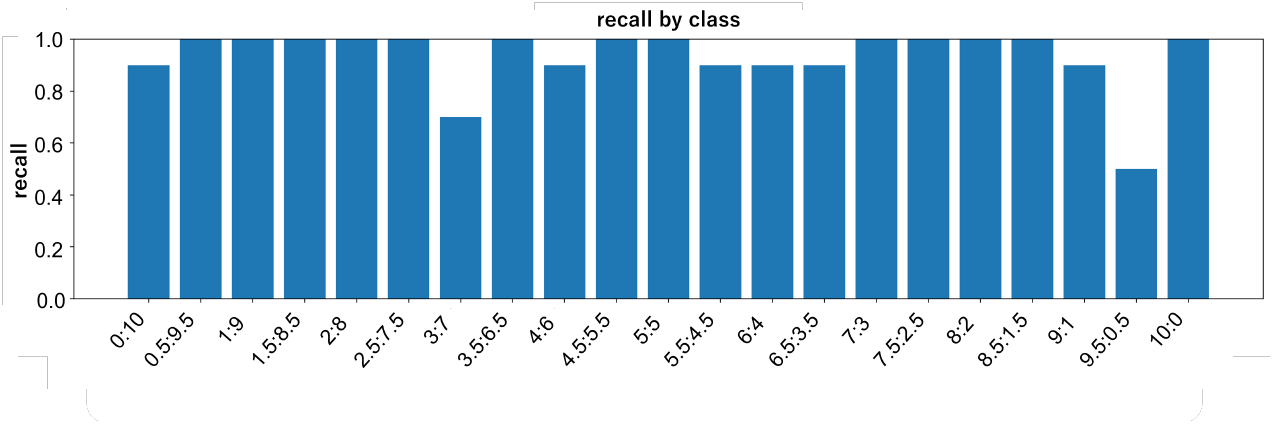
Class-wise recall of mixture-ratio classification using the 24 motion features with TCN.

**Fig 13.**
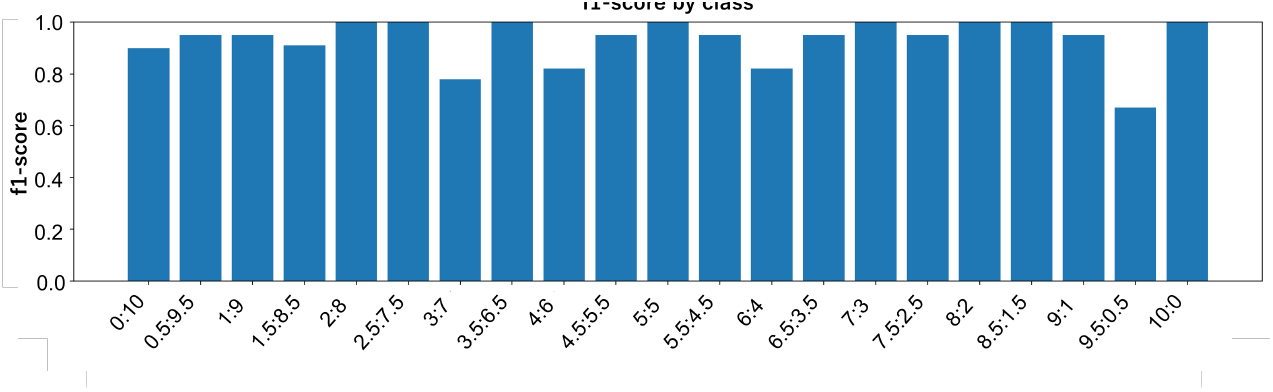
Class-wise F1 score of mixture-ratio classification using the 24 motion features with TCN.

Most mixture-ratio classes exhibited consistently high values for all three metrics, indicating stable classification performance without strong bias toward particular classes. In many intermediate mixture conditions, both precision and recall reached values close to 0.9, suggesting that the model effectively suppressed both false positives and false negatives. The correspondingly high F1 scores confirm that this balance was maintained simultaneously, rather than being achieved at the expense of one metric over the other. However, several classes showed relatively lower recall and F1 scores. This behavior can be attributed to the similarity of temporal motion patterns between neighboring mixture ratios. Because the TCN [18] learns temporal dependencies across the entire sequence, the boundaries between adjacent mixture conditions can become ambiguous when their motion dynamics are similar. In particular, classes near the 5:5 ratio—where *V. harveyi* and environmental bacteria coexist in comparable proportions—exhibited slightly reduced recall, suggesting that intermediate compositions are inherently harder to discriminate than extreme ratios dominated by a single species. In addition, *harveyi* exhibits temporally variable motility, including intermittent bursts of active motion and transient stationary periods. Such local fluctuations may influence the temporal feature representation learned by the model, occasionally leading to misclassification into neighboring mixture classes.

Overall, the results demonstrate that the proposed 24-feature representation combined with TCN [18] provides highly accurate classification performance. This indicates that temporal patterns of bacterial collective motion contain sufficient information to distinguish different mixture ratios with high reliability.

### Comparison with Alternative Models

To evaluate the effectiveness of the proposed TCN [18], its performance was compared with several baseline models, including XGBoost [29], SVM [27], and 1D-CNN [28]. The classification accuracies of the models are summarized in Table 1. Among the evaluated models, the proposed TCN [18] achieved the highest accuracy of 0.933. XGBoost [29] obtained an accuracy of 0.857, followed by SVM [27] (0.848) and 1D-CNN [28] (0.842).

**Table 1.** Classification accuracy of different models for mixture-ratio prediction.

| Model | Accuracy |
| --- | --- |
| TCN | **0.933** |
| XGBoost | 0.857 |
| SVM | 0.848 |
| 1D-CNN | 0.842 |

As shown in Fig 10, the TCN [18] model correctly classifies most mixture-ratio conditions, with predictions concentrated along the diagonal of the confusion matrix, indicating that the proposed model can reliably distinguish different bacterial mixture ratios. Confusion matrices of the baseline models are provided in the Supporting Information (Figs. S1–S3). Compared with TCN [18], the baseline models show increased misclassifications, particularly between neighboring mixture ratios, suggesting that models without explicit temporal modeling have difficulty capturing subtle differences in motion dynamics. While XGBoost [29] and SVM [27] treat each sample as a static feature vector and cannot fully utilize temporal structure, the 1D-CNN [28] model captures local temporal patterns but has a limited receptive field.

Overall, the superior performance of the TCN [18] suggests that explicit modeling of temporal structures is essential for accurately distinguishing mixture ratios from motion features. These results support the hypothesis that temporal patterns of bacterial collective motion contain discriminative information that cannot be fully captured by static feature models.

### Feature Importance Reveals possible Biological Mechanisms

To investigate which motion features contributed most to mixture-ratio classification, a feature importance analysis was conducted using permutation importance. The importance scores of all 24 motion features are shown in Fig 14.

**Fig 14.**
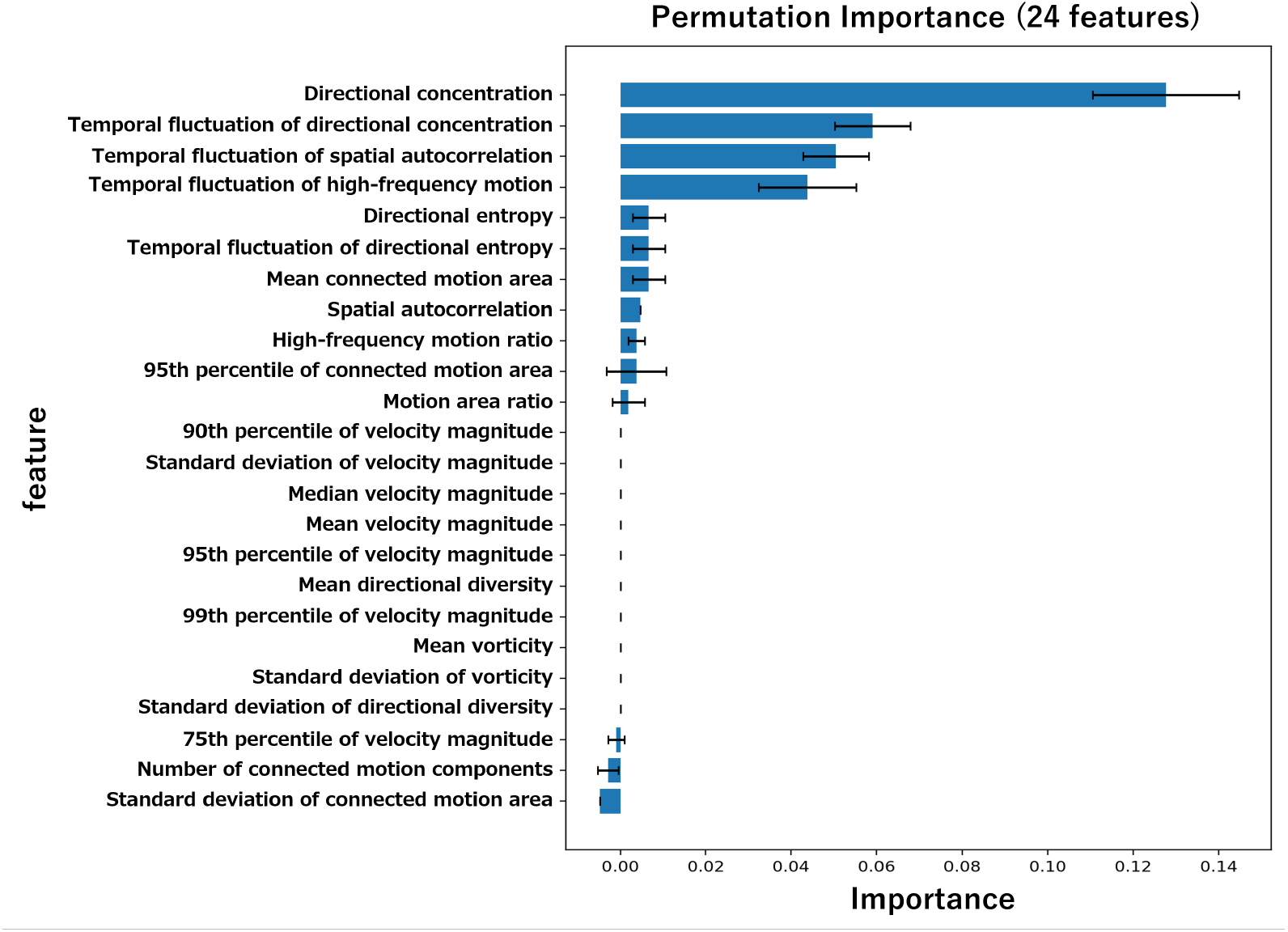
Permutation importance of the 24 motion features for mixture-ratio classification using TCN.

Among the evaluated features, directional concentration showed the highest importance score. Directional concentration is a metric that quantifies the degree of alignment in the movement directions of a bacterial population, measuring how uniformly the individual motion directions are oriented. The outstanding importance of this feature suggests that, in distinguishing mixture ratios, the collective directionality and the presence or absence of oriented motion across the bacterial population serve as more dominant discriminative criteria than the absolute magnitude of displacement or velocity.

In addition, several features representing temporal variability of motion patterns also exhibited relatively high importance scores. Temporal fluctuation of directional concentration quantifies the temporal variability of alignment, capturing how much the degree of directional coherence within the bacterial population changes over time. Temporal fluctuation of spatial autocorrelation represents the temporal variability of spatial coherence, reflecting how stably the spatial consistency of motion patterns is maintained across time. Temporal fluctuation of high-frequency motion represents the temporal variability of the high-frequency ratio, capturing how much the proportion of high-frequency components in the motion fluctuates over time. These features collectively serve as indicators of short-term fluctuations and instability in motion characteristics, reflecting whether bacterial motion is temporally stable or irregularly varying. Since TCN [18] is a model capable of explicitly learning temporal structures, it is considered that these temporally varying features were effectively utilized for classification. In contrast, features derived from simple velocity statistics, such as the mean velocity magnitude, the standard deviation of velocity magnitude, and other velocity quantile features, exhibited very low importance scores. This indicates that the overall magnitude of motion provides limited discriminative information for distinguishing mixture ratios in the present dataset. Furthermore, some features exhibited negative importance scores. A negative importance score indicates that including the feature degrades classification accuracy, implying that such features function as noise and adversely affect the discriminative performance of the model.

Overall, the results shown in Fig 14 suggest that mixture-ratio classification is primarily determined by the structure of collective motion rather than by simple activity intensity. In particular, directional alignment and temporal variability of motion patterns play key roles in characterizing bacterial population dynamics under different mixture conditions. These findings suggest that the collective organization of bacterial motion encodes richer information about population composition than simple measures of motility strength.

### Reduced Feature Model

To examine how classification performance changes with a reduced number of motion features, experiments were conducted using the most important feature and the four most important features identified by permutation importance. The confusion matrices for these reduced feature models are provided in the Supporting Information (Figs. S4 and S5). Using only the most important feature, directional concentration, the classification accuracy was 0.591. When the top four features were used, the accuracy improved to 0.657. A quantitative comparison of classification performance is summarized in Table 2. These results indicate that key characteristics of bacterial collective motion can be captured by a small number of features, while integrating multiple complementary descriptors improves classification performance. In particular, directional organization of bacterial motion provides primary discriminative information, and additional features refine classification by capturing subtle differences between similar mixture conditions. Reducing the number of required features while maintaining reasonable performance is important for practical applications, as it reduces computational cost and facilitates real-time implementation in automated screening systems.

**Table 2.** Classification accuracy of TCN using different feature sets.

| Case | Features Used | Accuracy |
| --- | --- | --- |
| All features | All 24 motion features | 0.933 |
| Top 4 features | Directional concentration,<br>Temporal fluctuation of directional alignment,<br>Temporal fluctuation of spatial autocorrelation,<br>Temporal fluctuation of high-frequency motion | 0.657 |
| Top 1 feature | Directional concentration | 0.591 |

### Regression Perspective

To further investigate the relationship between motion features and mixture ratios, a regression-based analysis was conducted. The predicted values generally follow the overall trend of the true mixture ratios, indicating that the model captures the correspondence between motion features and mixture ratios. Quantitatively, the regression achieved an *R*^2^ score of 0.807 and a mean squared error of approximately 8.62%, demonstrating that the model can represent the global structure of mixture-ratio variation. However, deviations from the ideal relationship are observed, particularly in intermediate mixture conditions, indicating limited precision in distinguishing similar motion patterns. Compared with regression, the classification framework provides more stable and consistent discrimination, suggesting that classification is more suitable for mixture-ratio analysis in this study. Nevertheless, the regression results confirm that motion features contain continuous information about population composition, supporting the validity of the proposed feature representation.

## Discussion & Conclusion

In this study, we demonstrated that a TCN based on optical flow can accurately classify the abundance ratio of mixed bacterial populations, a task that is virtually impossible using static bright-field images. The combination of optical flow and TCN proved highly suitable for this approach. Optical flow captures dense, pixel-level spatiotemporal variations in velocity fields without the heavy computational burden of tracking individual cells, while the TCN effectively learns the long-range temporal dependencies and hierarchical structures inherent in these continuous motion features.

Permutation importance analysis revealed that the “collectiveness” of motion, specifically directional concentration, made the highest contribution to the classification performance. It is widely known in the field of active matter physics that bacterial populations can give rise to emergent collective behaviors—such as swarms, vortices, and active turbulence—driven by hydrodynamic interactions and steric collisions, particularly at high cell densities or near phase interfaces. Although careful verification is required to determine the exact physical relationship between our classification metrics and the well-documented bacterial collective motion, it is highly intriguing that our model identified macroscopic collective alignment as a key discriminator. Furthermore, features related to temporal fluctuations in bacterial motion were ranked among the most important contributors. This result likely reflects the highly motile and temporally variable swimming behavior characteristic of Vibrio species. In this pilot study, we validated the classification of specific pathogenic bacteria against a heterogeneous and undefined mixture of environmental bacteria. To clarify the biological significance of these motion metrics and their direct relationship to actual bacterial kinematics, systematic investigations using defined bacterial mixtures with known motility profiles are necessary. This remains a fascinating theme for future research.

Regardless, the present workflow, which can classify the characteristics of bacterial populations directly from bright-field time-lapse images, holds great promise for various applications. Beyond the simple detection of specific microorganisms, such as pathogens, this approach could be applied to evaluate drug responses and antibiotic resistance using motility indicators, or to screen bacteria using microcompartments, such as microchambers or droplets. Because this approach relies on standard bright-field microscopy and imposes a much lower computational burden than single-cell tracking, it demonstrates significant potential, especially in low-resource environments, as it circumvents the need for expensive analytical equipment or complex molecular techniques such as fluorescent labeling or genetic manipulation. Moreover, this framework is not limited to swimming motility; it can handle any biological features that exhibit temporal changes. For example, it could be applied to monitor critical microbial indicators, such as cell proliferation dynamics or the degradation of microscopic substrates (e.g., cellulose), significantly broadening its utility.

As this study serves as a pilot validation of the combination of optical flow and TCN, continuous and detailed verification is required to determine its universal applicability for identifying or classifying a broader range of microorganisms or their communities. In particular, the effects of cell density, imaging conditions, time-lapse intervals, and frame rates on the classification accuracy must be rigorously evaluated in future studies. Nevertheless, because optical flow can capture a diverse array of kinematic and temporal features, it is highly likely to accurately capture subtle, non-obvious dynamic signatures that might be missed by human observers or by conventional methods that rely on predefined, static evaluation axes. This study represents the first attempt to classify using machine learning the temporal dynamics of microbial population images, and we anticipate significant developments and broad applications based on this label-free, time-resolved framework in the future.

## Supporting information

**S1 Fig. Confusion matrix of XGBoost**. Confusion matrix obtained using the XGBoost model. Compared with TCN, more misclassifications are observed, particularly between neighboring mixture ratios.

**S2 Fig. Confusion matrix of SVM**. Confusion matrix obtained using the SVM model. The results show increased confusion among adjacent mixture classes compared to the proposed method.

**S3 Fig. Confusion matrix of 1D-CNN**. Confusion matrix obtained using the 1D-CNN model. The model captures local temporal patterns but shows reduced performance compared with TCN.

**S4 Fig. Confusion matrix using the most important feature**. Confusion matrix obtained using only the directional concentration feature. The results show limited classification performance with substantial misclassifications across mixture ratios.

**S5 Fig. Confusion matrix using the top four features**. Confusion matrix obtained using the four most important features identified by permutation importance. Compared with the single-feature model, predictions are more concentrated around the diagonal.

**S6 Fig. Prediction versus true mixture ratios in the regression model**. Scatter plot of predicted versus true mixture ratios. The regression model captures the overall trend but shows deviations from the ideal diagonal line.

**S1 Table. List of motion features used in this study**. Summary of the 24 motion features, including descriptors related to velocity, directional alignment, spatial organization, and temporal variability.

**S1 Text. Detailed description of motion feature extraction**. Detailed definitions and computational procedures for all motion features.

**S1 Appendix. Model architecture and training details**. Additional details on the TCN model [18], hyperparameters, and training procedures are provided below. All experiments were implemented in Python using PyTorch [30], and optical flow was computed using the Farnebäck method [21] implemented in OpenCV [31]. Optimization was performed using the AdamW optimizer [32]. ReLU activation functions [33] were used throughout the network.

## Acknowledgments

This work was supported by the Japan Keirin Association (JKA) through its Promotion of Keirin (Grant numbers: 2023M-385 and 2025M-484).

